# Investigation of Bactericidal effect of Earwax on *Escherichia coli* and *Staphylococcus aureus* isolated from skin and stool samples of undergraduate students, Federal University of Agriculture Makurdi, Benue State, Nigeria

**DOI:** 10.1101/2024.07.29.605580

**Authors:** Aondongu Joseph Tsenongu, Tersagh Smart Ichor, Saaondo James Ashar, Ajekwe Stephen Mena, Avalumun Kwaghve

## Abstract

Earwax is a normal product of the ear, which protects the skin of the ear from water, dirt dust and infection. The aim of this study was to determine the antibacterial activity of earwax on *Staphylococcus aureus* and *Escherichia coli* isolated from skin and stool respectively. Bacteriological investigations were carried out using culture and morphological characteristics, conventional biochemical tests and antibiotic susceptibility testing of both earwax and conventional antibiotics against the presumptive isolates. Occurrence of *Staphylococcus aureus* (58.3%) and *Escherichia coli* (45.8%) in skin and stool samples were respectively determined based on cultural and morphological characteristics. Conventional biochemical tests were carried out on the 14 and 10 presumptive *Staphylococcus aureus* and *Escherichia coli* isolates and then 10 and 8 presumptive isolates were detected respectively. The antibacterial activity of the test agent was determined using the agar well diffusion technique; five different concentrations (20% v/v, 40% v/v, 60% v/v, 80% v/v, and 100% v/v) of the test agents were evaluated against the bacterial isolates. The earwax showed antibacterial activity against both isolates (except the 20% v/v for *Staphylococcus aureus*) at all concentrations used with antibacterial activity directly proportional to earwax concentration and a better antibacterial activity against *Staphylococcus aureus* (zone of inhibition 20mm) *aureus* than *Escherichia coli*. Antibiotic susceptibility testing was carried out on the 10(41.7%) and 8(33.3%) presumptive isolates of *Staphylococcus aureus* and *Escherichia coli* respectively based on Kirby Bauer method. Highest rate of susceptibilities of the *Staphylococcus aureus* isolates to antibiotics were observed in Erythromycin (100%), while for *Escherichia coli* were observed in Erythromycin and Levofloxacin (100%). Meanwhile, highest rate of resistant to *Staphylococcus aureus* isolates were observed in Gentamicin (100%) while for *Escherichia coli* was observed in Amoxicillin (100%). This study shows that earwax plays a critical role in the fight against infectious microorganisms as some of the antibiotics.

## Introduction

Earwax (cerumen) is a grey brown or yellowish waxy substance secreted in the outer one third of ear canal and, is a protective barrier between the external environment and deep external auditory canal (Wayne *et al*., 2020). Chemical composition of earwax includes 60% desquamated, 12-20% saturated and unsaturated fatty acid and 6-10% cholesterol (Yang *et al*., 2019). It is a secretion of specialized sets of glands, like sebaceous glands that secrete sebum (combination of fatty acid). Another gland i.e. apocrine sweat glands release secretion that combines with the sebum to form cerumen. It picks up discarded cells, ear follicles and may contain dust or other debris, but the resulting compound forms earwax or cerumen. It is also known by the medical term cerumen, is a gray-orange or yellowish waxy substance secreted in the ear canal of humans and other mammals. It protects the skin of human ear canal, assists in cleaning and also provides some protection against bacteria, fungi, insects, dust particles and water. Its antimicrobial activity is due to the presence of saturated fatty acids, lysozyme and slight acidity with the PH value of 6.1(Abimbola, 2019).

Earwax is a mixture of layer of keratinocytes cell which migrate from the deeper layer of the epidermis and finally shed from the skin and are secreted into the outer third of the wall of external auditory canal. This forms a greyish-black coloured thick substance which gets deposited inside the external auditory canal. Secretions from glands and cells from the hair in the canal also mix and forms earwax (Devi *et al.,* 2015). Earwax is composed of peeling sheets of corneocytes, originating from the deep and superficial external auditory canal, mixed with glandular secretions. It is a mixture of secretions from sebeceous glands modified apocrine sweat glands (Gupta *et al*., 2012). Cerumeneous glands in the auditory canal secrete lipids and peptides, respectively. Hairs in the external third of the canal also produce glandular secretions that contribute to earwax compositions (Gabriel, 2015). The wax produced forms a physiological barrier between the external environment and deep auditory canal. The chemical composition of the wax has also been believed to have antibacterial and anti fungal properties. It holds that wax contains antimicrobial properties which prevents external ear from infections (Sumit *et al., 2012*). Previous Studies conducted up until 1960s found little evidence supporting antibacterial activity for earwax (Nichols and Perry,1956), more recent studies have found that cerumen has a bactericidal effects on some strains of bacteria. cerumen has been found to reduce the viability of wide range of bacteria, including *haemophilus influenzae*, sometimes by as much as 99 % (Chai *et al*.,1980). The growth of two fungi commonly present in Otomycosis was also significantly inhibited by human earwax (Megarry *et al*.,1980). These antimicrobial properties are due principally to the presence of saturated fatty acids, lysozyme and especially to its slight acidity (pH typically around 6.1 in normal individuals). Conversely, others research has that earwax can support microbial growth and some earwax samples were found to have bacterial counts as high as 10g (Adegbiji *et al*., 2017).

Ear infection in any form has a varied microbial etiology, which influences the selection of an efficacious of anti-microbial agents. According to WHO survey, 42 million people worldwide have hearing loss where major cause is otitis media. Infection of the ear can be classified depending upon the site: otitis externa (infection of external ear) and otitis media (infection on middle ear). (Horton *et al*., 2020). Otitis is a bacterial infection of the ear canal caused by rupture in the normal skin or cerumen, which is a protective barrier in the presence of elevated humidity and temperature. It is commonly known as *swimmer’s ear*, though anything that disrupts this protective lipid layer can lead to the introduction and proliferation of bacteria. Trauma from cleaning the ears with fingernails or cotton buds has been identified as the most common predisposing factor locally (Harith *et al*., 2018).

However, in as much as, the role of human cerumen has been known to protect the external ear canal against infections; there are still agitations as regarding this topic. Few authors have suggested that earwax lacks the ability to prevent infections and that the rich nutrients of cerumen support abundant growth of bacteria and fungi. Moreover, the two most common bacterial species isolated from the external auditory canal of normal individuals are the *Staphylococcus* species (*Staphylococcus auricularis, Staphylococcus epidermidis, Staphylococcus aureus* and *Staphylococcus capitis*) and the *Corynebacterium* species (*Turicella otitidis* and *Corynebacterium auris*). The third most frequently recovered bacteria are the *Streptococci* and *Enterococci*. Together, they account for more than 90% of the normal flora in the external auditory canal. *Pseudomonas aeruginosa, S. epidermidis* and *S. aureus* are the most common pathogenic bacteria isolated in acute diffuse otitis external locally (Gerchman *et al.,* 2012).

## Materials and Methods

### Study Area

The research study was carried out at the Department of Microbiology, Federal University of Agriculture Makurdi, Benue State.

### Study Location

It is the capital of Benue State, located in the central Nigeria. The city is situated on the south bank of the Benue River. In 2016, Makurdi and the surrounding areas had an estimated population of 365,000 (Government of Benue State, 2020). It is predominantly an agricultural area specializing in cash crops and subsistence crops. Makurdi is located within latitudes 7.74° 7̍ꞌ, 8.51° 12^’^ N and longitudes 07° 41^’^ E in the Northern Guinea Savannah zone. The region is a tropical area with alternating wet (April to October) and dry (November to March) seasons and an annual average precipitation of 1240-1440 mm

### Equipment

Whatman filter papers, punching machine, sterile swab sticks, Petri dishes, sterile syringes, Conical flask, Test tubes, Incubator, Filter paper, Wire Loop, Cotton wool and Bijou bottle.

### Reagents

Sodium bicarbonate-glycerol buffer (pH 8.2), NaHCO3 (5%), Gram stain, peptone water and normal saline.

### Media

Mannitol Salt Agar (MSA), Eosin Methylene Blue Agar (EMBA) and Muller Hinton Agar (MHA), MacConkey broth and Nutrient agar.

### Sample Collection

After taken informed consent, 24 samples were collected each for skin and stool samples from undergraduate students of the department of Microbiology, Federal University of Agriculture, Makurdi using sterile swabs and bottles under proper hygienic conditions and then, they were transferred to the laboratory. Skin samples taken by scrubbing sterile swab sticks on the skin were directly dissolved in normal saline. Stool samples were inoculated in MacConkey broth for enrichment. Collected samples were suitably diluted and were streaked on nutrient agar medium. After incubation, these were subjected for isolation of *Escherichia coli* and *Staphylococcus aureus*.

### Cerumen Collection

Earwax was collected from 8 apparently healthy persons of both genders, ranging from 20 to 28 years using sterile swab sticks and the amount of cerumen from each was measured. Then, the samples pooled were dissolved in sodium bicarbonate-glycerol buffer (pH 8.2) (Hasibi *et al*., 2017).

### Isolation and Identification of *Escherichia coli*

Stool samples were processed to isolate *E. coli* as described by the Bacteriological Analytical Manual. The samples were inoculated into MacCkonkey broth for enrichment at 37°C for 24hrs. Afterwards, the 24 hours old pink coloured colonies were subcultured on Eosin methylene blue (EMB) agar by streak plate method. Colonies producing greenish metallic sheen on EMB Agar were suspected as having *E.coli.* The colonies that produced greenish metallic sheen colour were identified after various biochemical test as E. *coli* (Gerchman *et al*., 2012).

### Isolation and Identification of *Staphylococcus aureus*

Skin samples were dissolved in normal saline, streaked on Mannitol Salt Agar (MSA), incubated at 37°C for 24 hours. This grew and formed colonies with yellow colour. Significant colonies were picked up for Gram’s staining and biochemical tests were done for confirmation of *S.aureus.* The identification of *S.aureus* isolate was done based on colony morphology and biochemical tests like Indole, Catalase, and citrate utilization test. (Dubey *et al*., 2013).

### Phenotypic identification of isolates

The identification was done from microscopic observations, i.e. morphology and arrangement of cells, cultural characteristics on agar and in broth and from biochemical tests (Gupta *et al*., 2012). Loop full of suspected *E. coli* and *S. aureus* colonies were taken and streaked on nutrient agar and incubated for 24hours to observe the isolated colonies. Then all the plates were incubated at 37oC for 24-48hrs in an inverted position (Ambika *et al.,* 2015) to determine the specific cultural characters of isolates. Various biochemical tests were performed in order to identify and characterize the bacteria. The tests were performed according to the methods described by Gerchman *et al*. (2012).The tests include Gram staining, motility, catalase and indole.

### Biochemical identification

#### Catalase

Culture from a typical colony was placed on to a clean grease-free slide and a drop of 3% hydrogen peroxide (H_2_O_2_) solution was added on to the culture and closely observed for evolution of bubbles. The production of bubbles indicates positive catalase reaction and was recorded accordingly for the presence of or absence of enzyme (Gerchman *et al*., 2012).

#### Indole Test

To detect the ability of an organism to break tryptophan to pyruvate and indole. Most strain of Escherichia coli breakdown the amino acid and tryptophan with the release of indole. The test organism was inoculated into 5 ml of sterile peptone water in a bijou bottle and at 37 °C for 24 hours. Indole production was tested by the addition of 0.5 ml Kovac’s reagent to both culture, the development of bright pink color on the top layer of broth within 10-30 seconds indicated a positive result, while yellow color at the top of the broth indicated negative result (Gupta *et al*., 2012).

#### Motility Test

The soft agar stabbing tube method was done using nutrient agar. Nutrient agar slant was prepared and a sterile needle was used to stab the agar slant and incubated for 24hours at 37°C. Positive results shows well dispersed growth from line of inoculation (Horton *et al*., 2020).

#### Citrase Utilization Test

The isolates were inoculated in Simmons citrate agar incubated at 37°C for 24hours. After incubation, the appearance of blue coloration indicates the positive test for citrate utilization and was recorded accordingly for the isolates tested while green coloration indicated negative result (Gupta *et al*., 2012).

#### Gram Staining

Bacteria smear were prepared on a clean grease free microscopic slide and fixed with air and heat. Samples were then stained with crystal violet for 30 seconds and rinsed with water. After that samples were covered with gram iodine and allowed to act for 30 seconds and rinsed further with distilled water. A drop of95% alcohol was added and kept for 10-20 seconds for discoloration. The slides were further rinsed with sterile water. Counter staining was done with safranin for 20-30 seconds and slides were rinsed with sterile water and then dried (Vincent and Humphrey, 1970). Oil immersion was then dropped on the stained slides and was examined under compound microscope. This was examined using 100X objective lenses of compound microscope (Gerchman *et al*., 2012).

#### Antibiotic susceptibility testing

The antimicrobial susceptibilities of *Salmonella* isolates were conducted using a modified Kirby-Bauer disk diffusion method (Bauer *et al*., 1966). A panel of 10 antibiotics commonly used in human medicine were screened for Streptomycin (30µg), Gentamycin (10µg), Ciproflox (10µg), Ampiclox (10µg), Rocephin (5µg), Amoxacillin (30µg), Norbactin (10µg), Chloramphenicol (30µg), Erythromycin (30µg) and Levofloxacin (5µg) were tested. The surface of each inoculum was touched with a sterile inoculating loop and transferred into 4ml of Mueller Hinton broth. The turbidity of the inoculum was adjusted using sterile saline solution to a 0.5 McFarland standard. The diluted bacterial suspension was streaked onto Mueller Hinton agar plates using sterile cotton swabs. The plates were seeded uniformly by rubbing the swabs against the entire agar surface and allowed to dry for about 10 minutes. Each antimicrobial impregnated disk was applied onto the surface of the inoculated plate by using a disk dispenser (Oxoid^TM^). All plates were incubated at 37^0^C for 18-24 hours. Growth inhibition zones and classification of isolates as susceptible, intermediate and resistant was done according to recommendations of Clinical and Laboratory Standards Institute (CLSI, 2020). A reference strain of *E. coli* ATCC 25922 was used as quality control. Multidrug resistance (MDR) was defined as resistance of the isolate to at least one antibiotics in three or more classes of antibiotics (Magiorakos *et al.,* 2012). Multiple antibiotic resistance (MAR) index for each of the isolates was calculated by the formula given by Krumperman, (1983).

### Determination of Bactericidal Activity of Earwax

#### Cerumen Disc Method

To determine suitable concentration for the inhibition of *S.aureus* and *E.coli*, cerumen disc method was opted. Cerumen was weighed as 0.1g, 0.2g, and 0.3g, amounts and then dissolved in 100ml each in sterile buffer (pH 8.2) mixture of NaHCO3 (5%) and glycerol (30%).The cerumen-buffer mixture was homogenized by repeated passage through a series of needles ranging from 19 to 23guage with sterile syringes. This procedure broke the cerumen into fine particles distributed evenly in buffer and resulted in a milky suspension. Whatman filter papers were punched with a punching machine as small discs and were sterilized. Using sterile syringes 1ml of each isolates was inoculated in small discs on solidified Muller Hinton Agar plates containing 1ml of earwax concentrations each. These plates were labelled as corresponding to the cerumen (Vaghela *et al*., 2023). Then all the plates were incubated at 37°C for 48hours. After incubation, zones of inhibition were measured, results were recorded and also this was compared with suitable antibiotic concentration, which was inhibitory against the same bacteria (CLSI, 2017).

## Results

In this study, the occurrence of *Staphylococcus aureus* (58.3%) and *Escherichia coli* (45.8%) in skin and stool samples were respectively determined based on cultural and morphological characteristics (Table 1).

**Table 1:** Occurrence of *Staphylococcus aureus* and *Escherichia coli* in Skin and Stool of undergraduate students, Department of Microbiology, Federal University of Agriculture, Makurdi based on cultural and morphological characteristics.

| Type of samples | Total number of samples collected | Number of positive | % of positive | Presumptive inference |
| --- | --- | --- | --- | --- |
| Skin | 24 | 14 | 58.3 | *Staphylococcus aureus* |
| Stool | 24 | 11 | 45.8 | *Escherichia coli* |

Conventional biochemical tests were carried out on the 14 presumptive *Staphylococcus aureus* isolates and 10 of *Escherichia coli,* and 10 and 8 presumptive isolates were detected respectively (Table 2a and b).

**Table 2a:** Presumptive conventional biochemical tests of *Staphylococcus aureus* from undergraduate students, Department of Microbiology, Federal University of Agriculture, Makurdi.

| S/No | Sample No | Indole | Catalase | Citrate | Motility | Presumptive inference |
| --- | --- | --- | --- | --- | --- | --- |
| 1 | Sk1 | - | + | + | + | *Staphylococcus aureus* |
| 2 | Sk2 | + | - | - | + | Not applicable |
| 3 | Sk3 | + | + | - | + | Not applicable |
| 4 | Sk4 | - | + | + | + | *Staphylococcus aureus* |
| 5 | Sk5 | + | + | - | + | Not applicable |
| 6 | Sk6 | - | + | - | + | Not applicable |
| 7 | Sk7 | - | + | + | + | *Staphylococcus aureus* |
| 8 | Sk8 | + | - | - | + | Not applicable |
| 9 | Sk9 | + | - | - | + | Not applicable |
| 10 | Sk10 | + | - | - | + | Not applicable |
| 11 | Sk11 | - | + | + | + | *Staphylococcus aureus* |
| 12 | Sk12 | - | + | - | + | Not applicable |
| 13 | Sk13 | - | + | - | + | Not applicable |
| 14 | Sk14 | - | + | + | + | *Staphylococcus aureus* |
| 15 | Sk15 | - | + | - | + | Not applicable |
| 16 | Sk16 | + | - | - | + | Not applicable |
| 17 | Sk17 | - | + | + | + | *Staphylococcus aureus* |
| 18 | Sk18 | + | - | - | + | Not applicable |
| 19 | Sk19 | + | - | - | + | Not applicable |
| 20 | Sk20 | - | + | + | + | *Staphylococcus aureus* |
| 21 | Sk21 | - | + | + | + | *Staphylococcus aureus* |
| 22 | Sk22 | - | + | + | + | *Staphylococcus aureus* |
| 23 | Sk23 | - | + | + | + | *Staphylococcus aureus* |
| 24 | Sk24 | + | - | - | + | Not applicable |

(Sk = Skin, + = positive reaction, - = negative reaction)

**Table 2b:** Presumptive conventional biochemical tests of *Staphylococcus aureus* from undergraduate students, Department of Microbiology, Federal University of Agriculture, Makurdi.

| S/No | Sample No | Indole | Catalase | Citrate | Motility | Presumptive inference |
| --- | --- | --- | --- | --- | --- | --- |
| 1 | St1 | + | + | - | + | *Escherichia coli* |
| 2 | St2 | + | + | - | + | *Escherichia coli* |
| 3 | St3 | + | + | - | + | *Escherichia coli* |
| 4 | St4 | - | - | - | + | Not applicable |
| 5 | St5 | - | + | + | + | Not applicable |
| 6 | St6 | - | - | - | + | Not applicable |
| 7 | St7 | + | + | - | + | *Escherichia coli* |
| 8 | St8 | - | + | + | + | Not applicable |
| 9 | St9 | - | - | - | + | Not applicable |
| 10 | St10 | - | + | + | + | Not applicable |
| 11 | St11 | + | + | - | + | *Escherichia coli* |
| 12 | St12 | - | + | + | + | Not applicable |
| 13 | St13 | - | - | - | + | Not applicable |
| 14 | St14 | - | + | + | + | Not applicable |
| 15 | St15 | + | + | - | + | *Escherichia coli* |
| 16 | St16 | - | + | + | + | Not applicable |
| 17 | St17 | - | - | - | + | Not applicable |
| 18 | St18 | - | + | + | + | Not applicable |
| 19 | St19 | + | + | - | + | *Escherichia coli* |
| 20 | St20 | - | + | + | + | Not applicable |
| 21 | St21 | - | - | - | + | Not applicable |
| 22 | St22 | - | + | + | + | Not applicable |
| 23 | St23 | + | + | - | + | *Escherichia coli* |
| 24 | St24 | - | + | + | + | Not applicable |

(St = Stool, + = positive reaction, - = negative reaction)

The result of the antibacterial activity of the earwax against the *Staphylococcus aureus* and *Escherichia coli* were shown below (table 3). The earwax showed antibacterial effect on both isolates. The highest zone of inhibition was observed at the 100% v/v earwax concentration which was 18mm and 20mm for the *Staphylococcus aureus* and *Escherichia coli* respectively. There was an increasing antibacterial activity with increasing earwax concentration. No antibacterial activity was observed at the 20% v/v concentration of the earwax against *Staphylococcus aureus*.

**Table 3:** Antibacterial activity of Earwax against *Staphylococcus aureus* and *Escherichia coli*.

| Concentration (% vol/vol) | Diameter of Zone of Inhibition (mm) |  |
| --- | --- | --- |
|  | *Staphylococcus aureus* | *Escherichia coli* |
| 20 | - | 6 |
| 40 | 12 | 15 |
| 60 | 13 | 14 |
| 80 | 15 | 17 |
| 100 | 20 | 18 |

Key: - = no zone of inhibition
Control: distilled water (zone of inhibition = 0mm)

Antibiotic susceptibility testing was carried out on the ten and seven presumptive isolates of *Staphylococcus aureus* and *Escherichia coli* respectively based on Kirby Bauer method. Highest rate of susceptibilities of the *Staphylococcus aureus* isolates to antibiotics were observed in Erythromycin (100%), while for *Escherichia coli* were observed in Erythromycin and Levofloxacin (100%). Meanwhile, highest rate of resistant to *Staphylococcus aureus* isolates were observed in Gentamicin (100%) while for *Escherichia coli* was observed in Amoxicillin (100%) (Table 4a and b).

**Table 4a:** Antimicrobial susceptibilities of ten (10) presumptive *Staphylococcus aureus* isolates to 10 antibiotics.

| **Antimicrobial Agent** | **Disk Code** | **Disk content (µg)** | **No. (%) isolates Susceptible** | **No. (%) isolates Intermediate** | **No. (%) isolates Resistant** |
| --- | --- | --- | --- | --- | --- |
| Gentamicin | CN | 10 | 0(0) | 0(0) | 10(100) |
| Ciproflox 1 | CF | 10 | 5(50) | 0(0) | 5(50) |
| Ampiclox | AP | 10 | 4(40) | 2(20) | 4(40) |
| Amoxicillin | AMC | 10 | 0(0) | 1(10) | 9(90) |
| Rocephin | RC | 5 | 9(90) | 0(0) | 1(10) |
| Norbactin | NT | 10 | 3(30) | 4(40) | 3(30) |
| Chloramphenicol | C | 30 | 5(50) | 1(10) | 4(40) |
| Erythromycin | EC | 30 | 10(100) | 0(0) | 0(0) |
| Levofloxacin | LF | 5 | 4(40) | 2(20) | 4(40) |

**Table 4b:** Antimicrobial susceptibilities of seven (7) presumptive *Escherichia coli* isolates to 10 antibiotics.

| **Antimicrobial Agent** | **Disk Code** | **Disk content (µg)** | **No. (%) isolates Susceptible** | **No. (%) isolates Intermediate** | **No. (%) isolates Resistant** |
| --- | --- | --- | --- | --- | --- |
| Gentamicin | CN | 10 | 3(42.9) | 0(0) | 4(57.1) |
| Ciproflox 1 | CF | 10 | 5(71.4) | 0(0) | 2(28.6) |
| Ampiclox | AP | 10 | 4(57.1) | 2(28.6) | 1(14.3) |
| Amoxicillin | AMC | 10 | 0(0) | (0) | 7(100) |
| Rocephin | RC | 5 | 4(57.1) | 0(0) | 3(42.9) |
| Norbactin | NT | 10 | 1(14.3) | 2(28.6) | 3(42.9) |
| Chloramphenicol | C | 30 | 5(71.4) | 0(0) | 2(28.6) |
| Erythromycin | EC | 30 | 7(100) | 0(0) | 0(0) |
| Levofloxacin | LF | 5 | 7(0) | (0) | 0(0) |

## Discussion

This study demonstrated the occurrence of *Staphylococcus aureus* (58.3%) and *Escherichia coli* (45.8%) based on cultural and morphological characteristics. Meanwhile, conventional biochemical tests identified 10(41.7%) and 8(33.3%) presumptive isolates of *Staphylococcus aureus* and *Escherichia coli*. This is similar to work documented by Prasad and Moha, (2014) but contrary to work carried out by Shokry and Antoniosi-Filho, (2017), and this may be attributed to the type of population and sample size.

Variable results observed by different authors regarding antibacterial properties of human earwax. Most authors, however strongly postulated that human wax alone with it barrier host defense mechanism, also carry some amount of antibacterial effect which does inhibit growth of microorganisms (Shokry *et al*., 2017). Role of wax in creating a physical barrier between external and internal environment has also been given. When wax is removed this barrier will be lost, and results in bacterial growth leading to infections (Kilkenny, 2019). However, there has been no tangible evidence to support this view. The role of common resident flora in the region might also play a role which again questions the overall ability of human wax as an antbacterial agent. In our study, earwax was found to inhibit growth of *Staphylococcus aureus* and *Escherichia coli* at different concentrations which agreed with the findings of Nagaraj *et al. (*2019) and Zeeshan a *et al*. (2018). The study demonstrated that earwax has antibacterial property, which plays a role in protection of the external auditory canal against bacteria.

The result of the antibacterial activity of the test agent called earwax showed that the earwax had antibacterial effect on both isolates. The highest zone of inhibition was observed at the 100% v/v earwax concentration which was 18mm and 20mm for the *Staphylococcus aureus* and *Escherichia coli* respectively. There was an increasing antibacterial activity with increasing earwax concentration. No antibacterial activity was observed at the 20% v/v concentration of the earwax against *Staphylococcus aureus.*. In our findings, *Escherichia coli* was more susceptible to earwax as compared to *Staphylococcus aureus*. This implies that, it was the most non-resistant strain of *Escherichia coli* towards earwax. The significant bactericidal activity of cerumen on *Escherichia coli* might be as a result of not been a normal commensal of the ear as reported by Ngo *et al*. (2016). *Staphylococcus aureus* was resistant to earwax at 20% v/v, hence it showed no zone of inhibition, which might be due to variation in strains and different efficacy of earwax as collected from different individuals. The result corroborated with the findings of Saxby *et al*. (2012) who reported inconsistent bactericidal activity of earwax on *Staphylococcus aureus*.

The emergence of antimicrobial resistance is mainly promoted by the use of antibiotics in animal feed to promote the growth of food animals, and in veterinary medicine to treat bacterial infections in those animals while humans is to treat infections (Awanye *et al.,* 2022).Though, there is policy guiding the use of antimicrobials in humans and animals in Nigeria, there is high level of indiscriminate use of antimicrobials in among humans (Ojo *et al.,* 2012). The antibiotic susceptibilities of the isolates showed varying degrees of sensitivity. Greater proportions of the presumptive *Staphylococcus aureus* isolates were susceptible to Erythromycin (100%), while for *Escherichia coli* were observed in Erythromycin and Levofloxacin (100%). Meanwhile, highest rate of resistant to *Staphylococcus aureus* isolates were observed in Gentamicin (100%) while for *Escherichia coli* was observed in Amoxicillin (100%). This finding is similar to the work of Fagbamila*et al*. (2017) in Plateau who reported that the highest level of resistance (100%) of the isolates was to oxacillin, but contrary to that of Jasini *et al*. (2020) in Borno who reported 100% resistance of the *Salmonella* isolates to gentamicin. This may be as result of the choice and level of usage of the antibiotics in various locations. This suggests that antibiotics still play a critical role and are important in the fight against infectious microorganisms

## Conclusion

The study showed significant role played by human earwax in the prevention of ear infections. The study divulges the ability of the earwax to inhibit growth of *Staphylococcus aureus* and *Escherichia coli* which are the known agents for a disease condition called Otitis externa. Present work found bactericidal activity of human wax to be more effective on *Escherichia coli.* This effectiveness of the earwax on *Escherichia coli* was due to the fact that, earwax posesses human antimicrobial protein known as B-Defensins which has strong effect on Gram-negative bacteria.

It was also seen that antibiotics still play a critical role and are important in the fight against infectious microorganisms as some of antibiotics such Erythromycin and Levofloxacin were able to inhibit the growth of *Staphylococcus aureus* and *Escherichia coli*.

This study justifies the necessity to avoid routine removal of earwax. The information derived from this work may be of immense value in education of the public on the importance of the human earwax in preventing ear infections and may also assist clinicians, physicians and public health workers in the rational choice of antibiotics therapy and reduce the misuse of antibiotics.

## Recommendations

Based on these findings, the following are recommended;

1. Routine removal of the cerumen is unnecessary except the impacted wax is leading to earache or conductive hearing loss
2. Public health workers, clinicians should educate the public on the role human earwax play against ear infections and when it becomes necessary to be removed.
3. Adequate training is reccomended for physicians and nurses on the proper approaches for removal of earwax with minimal risks

## References

Abimbola, O.(2019). A study on tongue rolling, tongue folding and cerumen type in a Nigeria population. Anatomy Journal of Africans, 7(8):15–40

Adegbiji, A., Aremu, S., and Olatoke, E. (2017). Epidemiology of otitis external in developing country. International Journal of Sciences, 8(1):18–23.

Awanye, A., Chidozie, N., Stanley, C., Onah, H., Okonko, I., Egbe, N. (2022). Multidrug-Resistant and Extremely Drug-Resistant *Pseudomonas aeruginosa* in Clinical Samples from a Tertiary Healthcare Facility in Nigeria. Journal of Microbiological Research,19(4):447–454

Bauer, B., Kirby, B., Varsha, G., Atul, S., Vishal, G., and Jagdish, C. (1966). Patterns in antimicrobial susceptibility of Salmonellae isolated at a tertiary care hospital in northern India. Journal of Microbiological Research,1(134):4–8

Clinical and Laboratory Standards Institute (CLSI). Performance standards for antimicrobial susceptibility testing. Clinical and Laboratory Standards Institute (2017).

Devi, K., Lakshmi, B., Ratnasri, P., and Hemalatha, K.(2015) Antimicrobial activity of cerumen. Journal of Microbiology Research, 5(3):670–80.

Fagbamila, I., Lisa, B., Marzia, M., Kwaga, J., Sati, S., Paola, Z., Antonia, A., Lettini, M., Paul, A., Junaidu. K., Umoh, J., Antonia, R., and Muhammad, M. (2017). *Salmonella* serovars and their distribution in Nigerian commercial chicken layer farms. Journal of Microbiological Research,9(2):47–54

Gabriel, O. (2015). Cerumen impaction: Challenges and management profile in a rural health facility. Nigerian Medical Journal, (5)6:3–90.

Gerchman, Y., Patichov, R., and Zeltzer, T. (2012). Lipolytic, proteolytic, and cholesteroldegrading bacteria from the human cerumen. Journal of Microbiology Research, 6(4):588–91.

Gupta, S., Singh, R., and Kosaraju, K.(2012). A study of antibacterial and antifungal properties of human cerumen. Indian Journal of Microbiology Research,11(6):85–811

Horton, G., Simpson, M., Beyea, M., and Beyea, A. (2020). Cerumen Management: An Updated Clinical Review and Evidence-Based Approach for Primary Care Physicians. Journal of Primary Care Community Health, 1(1):21–50

Harith, S., Mazlun, H., Mydin, M. (2018). Studies on phytochemical constituents and antimicrobial properties of Citrullus lanatus peels. Malaysian Journal Analytic Sciences, 22(6):151–156.

Hasibi, M., Ashtiani, K., and Motassadi-Zarandi, M. (2017). A treatment protocol for management of bacterial and fungal malignant external otitis: a large cohort in Tehran, Iran. Journal of Microbiology Research,12(6):561–567.

Jasini, A., Barka, J., and Auwalu, M. (2020). Phenotypic Identification and Antimicrobial Susceptibility Profile of *Salmonella* from Local and Exotic Chicken in Maiduguri, Nigeria. Journal of Microbiological Research, 4(5): 1816–4935

Kilkenny, N. (2019). The nurse’s role in earcare: undertaking hearing assessment and ear cleaning. British Journal of Nursing, 2(8):1–3.

Krumperman, P. H. (1983). Multiple antibiotics resistance indexing *Escherichia coli* to identify risk sources of faecal contamination of foods. Applied and Environmental Microbiology, 46: 165–170

Magnet, M., Arongozeb, M., Khan, M., Ahmed, A. (2013). Isolation and identification of different bacteria from different types of burn wound infections and study their antimicrobial sensitivity pattern. Journal of Microbiology Research, 13(1):125–32.

Nagaraj, G., Girdhar, A., and Chinnappa, J. (2019). Bacterial Profile of Middle Ear Fluid with Recurrent Acute Otitis Media Infection Using Culture Independent 16S rDNA Gene Sequencing. Journal of Pediatric Infectious Diseases, 14(8):108–15.

Ngo, C., Massa, H., Thornton, R., and Cripps, W. (2016). Predominant bacteria detected from the middle ear fluid of children experiencing otitis media: a systematic review. PLOS One Journay, 11(1):5–19.

Ojo, O. E., Ogunyinka, O. G., Agbaje, M., Okuboye, J. O., Kehinde, O. O. and Oyekunle, M. A. (2012). Antibiogram of *Enterobacteriaceae* isolated from free-range chickens in Abeokuta, Nigeria. VeterinaryArchives, 82: 577–589

Prasad, S., and Mohan, N.(2014). Anti-microbial properties of human wax. International Journal of Medical Sciences, 3(6):96–99

Saxby, C., Williams, R., and Hickey, S. (2012).Finding the most effective cerumenolytic. Journal of Laryngology, 12(7):1067–70.

Shareef, A., Amin, A., and Amin, B. (2018). Bacteriology of Otitis Media with Effusion in Children and Their Sensitivity to Antibiotics in Erbil City. Journal of Medical Sciences, 18(9):4–11

Shokry, E., and Antoniosi-Filho, R. (2017). Insights into cerumen and application in diagnostics: past, present and future prospective. Journal of Biochemical Medicine, 2(7):477–91.

Shokry, E., de Oliveira, A., and Avelino, M. (2017). Earwax: A neglected body secretion or a step ahead in clinical diagnosis? A pilot study. Journal of Proteomics, 1(5)9:92–101

Swain, K., Anand, N., and Sahu C.(2019). Human cerumen and its antimicrobial properties: Study at a tertiary care teaching hospital of Eastern India. Ann Indian Academy Otorhinolaryngol Head Neck Surgery. Indian Journal of Medicine, 3(5):13–16.

Swain, K., Sahu, C., Debta, P., and Baisakh, R.(2018). Antimicrobial properties of human cerumen. Journal of Applied Medicine, 1(8):15–19.

Tipton, B., Honsinger, L., Harris, S., and Malhotra, S. (2019). Acute Otologic Infections in Pediatric Patients. Journal of Pediatry Infectious Diseases,1(4):52–62

Vaghela, M., Doshi, H., and Rajput S. (2023) An analysis of ear discharge and antimicrobial sensitivity used in its treatment. International Journal of Medical Sciences, 4(5):2656–60.

Wayne, A., Parin, U., Erbas, G., and Ural. (2020). Investigation of bacterial and fungal agents from cutaneous lesions in canine Leishmaniosis. Indian Journal Animal Reserve, 5(4):96–100.

Yang, J., Chang, T., and Jiang, Y.(2018). Commensal Staphylococcus aureus provokes immunity to protect against skin infection of methicillin-resistant Staphylococcus aureus. International Journal of Molecular Sciences, 1(9):12–90.

Zeeshan, M., Zeb, J., and Saleem, M. (2018) ENT diseases presenting to a tertiary care hospital. Endocrinology Metabolism International Journal, 8(1):416–530

